# Quantitative evaluation of a high resolution lipidomics platform

**DOI:** 10.1101/627687

**Authors:** Juan Liu, Xiaojing Liu, Zhengtao Xiao, Jason W. Locasale

**Affiliations:** Department of Pharmacology and Cancer Biology, Duke University School of Medicine, Durham NC 27710

**Keywords:** Metabolomics, Lipidomics, Metformin, High Resolution Mass Spectrometry

## Abstract

Given the general importance of lipids in health and disease, there is a need for efficient technology that broadly profiles and quantitates the lipid composition of complex mixtures. In this study, we developed and quantitatively evaluated a platform that simultaneously profiles both lipids and polar metabolites from the same sample. This method was achieved by using a methyl tert-butyl ether (MTBE) extraction and employing two liquid chromatography methods coupled with high resolution mass spectrometry (LC-HRMS). This workflow enabled detection and semi-quantitation of over 300 polar metabolites as well as over 300 lipids with comprehensive coverage of diverse chemical classes. Using cultured mammalian cells as an example, we report the quantitative properties of the platform including the sensitivity and linear range. The lipidomics strategy was further applied to characterize changes to lipid metabolism upon treatment with metformin to human ovarian cancer cells. Of the 256 detected lipids, 99 lipids (39%) significantly increased, 11 lipids (4%) were significantly reduced and 146 lipids (57%) remain unchanged in metformin-treated cells. Stable isotope tracing of carbon into lipids using [^13^C_6_]-glucose further measured the contribution of *de novo* fatty acid synthesis to the total fatty acid pool. In summary, the platform enabled the semi-quantitative assessment of hundreds of lipid species and associated carbon incorporation from glucose in a high throughput manner.

## Background

Lipidomics, the simultaneous measurement of many lipid species, which has been propelled by analytical technologies based on mass spectrometry involving gas (GC) or liquid chromatography (LC) (1–4), is a powerful tool to understand metabolism (5–8). Gas chromatography coupled to mass spectrometry (GC-MS) has been a mainstay for lipidomics, which readily profiles volatile lipids and low molecular weight lipids (9). Liquid chromatography coupled with mass spectrometry (LC-MS) although lesser used, has shown promise in lipidomics with increased coverage of lipid classes and improved sensitivity (1, 10, 11). In addition to profiling of known compounds, LC coupled with high resolution mass spectrometry (HRMS) has facilitated the identification of new lipids (12).

Lipids are often extracted using chloroform and methyl tert-butyl ether (MTBE). MTBE is preferable for many reasons including its relative safety, and advantages such as ease of integration into automation for high-throughput, its low cost as well as its stability (13–15). In this study, we employed a combined methyl tert-butyl ether and our previously utilized methanol extraction method (13), which allows for the extraction and separation of lipids, polar metabolites, and proteins in a single extraction process, followed by metabolite analysis with hydrophilic interaction liquid chromatography (HILIC) and reversed-phase liquid chromatography (RPLC) coupled with HRMS. For this present study, we established a methodology using cultured mammalian cells, which resulted in the detection and semi-quantitation of over 300 polar metabolites as well as over 300 lipids with comprehensive coverage of diverse classes: free fatty acids, triacylglycerides, diacylglycerides, glycerophospholipids, sphingolipids, ceramides, sphingomyelins, and cholesterols. We further investigated the quantitative properties of the method including sensitivity and linear range of the lipodomics aspect of the method. The polar metabolomics method has been well characterized in our previous report (16). The lipidomics method developed in this report, allows semi-quantification over a linear range of 3 orders of magnitude and a sensitivity down to 10^4^ cells.

Furthermore as a proof of concept, we applied our platform to profile the lipid compositions of cultured ovarian cancer cells in response to metformin treatment, a common agent for the management of type II diabetes which has interesting anti-cancer properties (17, 18). Since this workflow measured both polar metabolites and lipids in one platform, we not only characterize the overall lipid profile, but also monitor the changes to polar metabolites related to lipid biosynthesis, such as glycerol-3-phopshate, the precursor for generating the back bone of the glycerol lipids. Following the development of this lipidomics analysis, we further combined the method with ^13^C tracing using ^13^C uniformly labeled glucose, considering glucose provides acetyl-CoA for *de novo* fatty acid synthesis but also contribute to glycerol-3-phosphate synthesis.

## Methods

### Materials

The human colon cancer cell line (HCT116) was obtained from ATCC. HeyA8 ovarian cancer (OvCa) cells were provided by Dr. Ernst Lengyel’s lab. Both human cell lines were validated using the cell line authentication service at Duke University and confirmed using the GenePrint 10 kit from Promega and tested to be mycoplasma-free. RPMI 1640 medium and dialyzed FBS were purchased from Life Technologies. Fetal Bovine Serum (FBS), penicillin, and streptomycin were purchased from Hyclone Laboratories. Optima grade ammonium hydroxide, acetonitrile, methanol, isopropanol (IPA) and water were purchased from Fisher Scientific. HPLC grade methyl tert-butyl ether was obtained from Alfa Aesar. HPLC grade acetic acid was obtained from Millipore.

### Cell culture

All cells were first cultured in a 10 cm dish with full growth medium (RPMI 1640 supplemented with 10 % FBS). The cell incubator was set at 37 °C in 5% CO_2_. For metabolite analysis, cells were seeded into 6 well plate at the density of 40 000 cells per well. After overnight incubation in full growth medium, for metformin treatment, the old medium was replaced with 1.5 ml of RPMI 1640 (supplemented with 10 % dialyzed FBS) with or without 1.5 mM metformin, incubated for 16h to reach equilibrium and replaced with the same medium following 48 h incubation. For ^13^C analysis, cancer cells were cultured in glucose-free RPMI medium containing [^13^C_6_]-glucose, supplemented with 10 % dialyzed FBS.

### Metabolite extraction

For lipid extraction, HeyA8 OvCa cells cultured in 6 well plate were rinsed with 0.1% NaCl and then extracted into a solution composed of methanol/MTBE/water (1:3:1, v/v). Samples were vortexed vigorously, followed by centrifugation at 3,500 g for 10 min at 5 °C. The top layer (lipids) and bottom layer (polar metabolites) were collected separately and dried in a vacuum concentrator. The dry pellets were stored at −80 °C until ready for LC-HRMS analysis. For dynamic range studies, HCT 116 cells were grown in a 10 cm cell culture dish to 80% confluency and followed the same extraction.

### LC-HRMS analysis

Lipid and polar metabolite extracts were reconstituted into sample solvent (50 μl isopropanol for lipids, 30 μl water:ACN:MeOH (2:1:1, v/v) for polar metabolites). The supernatant was transferred to LC vials for subsequent LC-HRMS analysis. An ultimate 3000 UHPLC is coupled to a Q Exactive Plus-mass spectrometer as previously described (16), with the metabolomics method documented in the same report. For lipidomics, MS parameters were the same as metabolomics, except a 4kV spray voltage with full scan at resolution of 70000 for both positive and negative modes with S-lens 65 were used. LC-HRMS raw data analysis was performed using Sieve 2.0 and integrated peak area was used to represent relative metabolite abundance. For lipids detected in both positive and negative ion modes, the ion with the higher intensity was used. A blank control following the same sample preparation using an empty tube was prepared and went through the same LC-HRMS analysis. Data from this blank control is used as a threshold to filter the background noise and artefactual ions. An integrated ion intensity of 1 × 10^5^ is defined to be lipid noise level. Lipids with poor signals (signal across all conditions were less than 1 × 10^5^) were filtered. For lipids with signal less than 1 × 10^5^ in one condition and signal more than 1 × 10^6^ in another condition, signal less than 1 × 10^5^ was replaced with 1 × 10^5^. See supplementary data for more details.

## Results

### Overview

First, we developed a platform that profiles a broad range of lipid classes (Figure 1). A cold MeOH/MTBE/H_2_O (1:3:1, v/v) extraction was used to achieve simultaneous extraction of both lipids (top organic layer) and polar metabolites (bottom aqueous layer). Polar metabolite profiling using HILIC-HRMS has been detailed in our previous report (16). Here, we focus on lipid profiling by applying RPLC-HRMS with positive and negative switching mode which broadens lipid coverage (Figure 1A). The reversed phase liquid chromatography method employing an Xbridge BEH C18 column (100 × 2.1 mm i.d., 2.5 μm; Waters) at 40°C is used for lipid separation. This column was selected for achieving a ~20 min running time to achieve high throughput. The mobile phase A is 80:20 (v/v) water:acetonitrile with 0.1% formic acid and 10 mM ammonium formate. The mobile phase B is 90:10 (v/v) isopropanol:acetonitrile with 0.1% formic acid and 10 mM ammonium formate. The linear gradient used is as follows: 0 min, 40% B; 1.5 min, 40% B, 5 min, 85% B; 12 min, 97% B, 16 min, 97% B, 16.5 min, 40% B, 20.5 min, 40% B. The flow rate is 0.2 mL/min. The covered lipid classes and elution order are demonstrated in Figure 1B for both positive mode (left panel) and negative mode (right panel). Figure 1C summarizes the detection of over 300 lipids (isomers are not differentiated and combined together as one lipid, represented by a combination of lipid class, number of carbons and number of C=C bonds in HCT116 cells, which include: 38 phosphatidylcholine diacyl (PC aa), 37 phosphatidylcholine acyl-ether (PC ae), 28 ceramides/dihydroceramides (Cer/ Cer(2H)), 27 phosphatidylethanolamine diacyl (PE aa), 19 triacylglycerides (TG), 18 phosphatidylethanolamine acyl-ethyl (PE ae), 17 free fatty acids, 16 2-hydroxyacyl-ceramides (Cer(2OH)), 16 2-hydroxyacyl-dihyroceramides (Cer((2H)/(2OH))), 16 diacylglycerides (DG), 12 lysophosphatidylcholine acyl (lyso PC a), 12 phosphatidylserine diacyl (PS aa), 8 sphingomyelin (SM), 7 phosphatidyl glycerol diacyl (PG aa), 7 lysophosphatidylethanolamine acyl (Lyso PE a), 7 monoacylglycerides (MG), 5 hydroxysphingomyelin (SM(OH)), and 5 cholesterol species.

**Figure 1.**
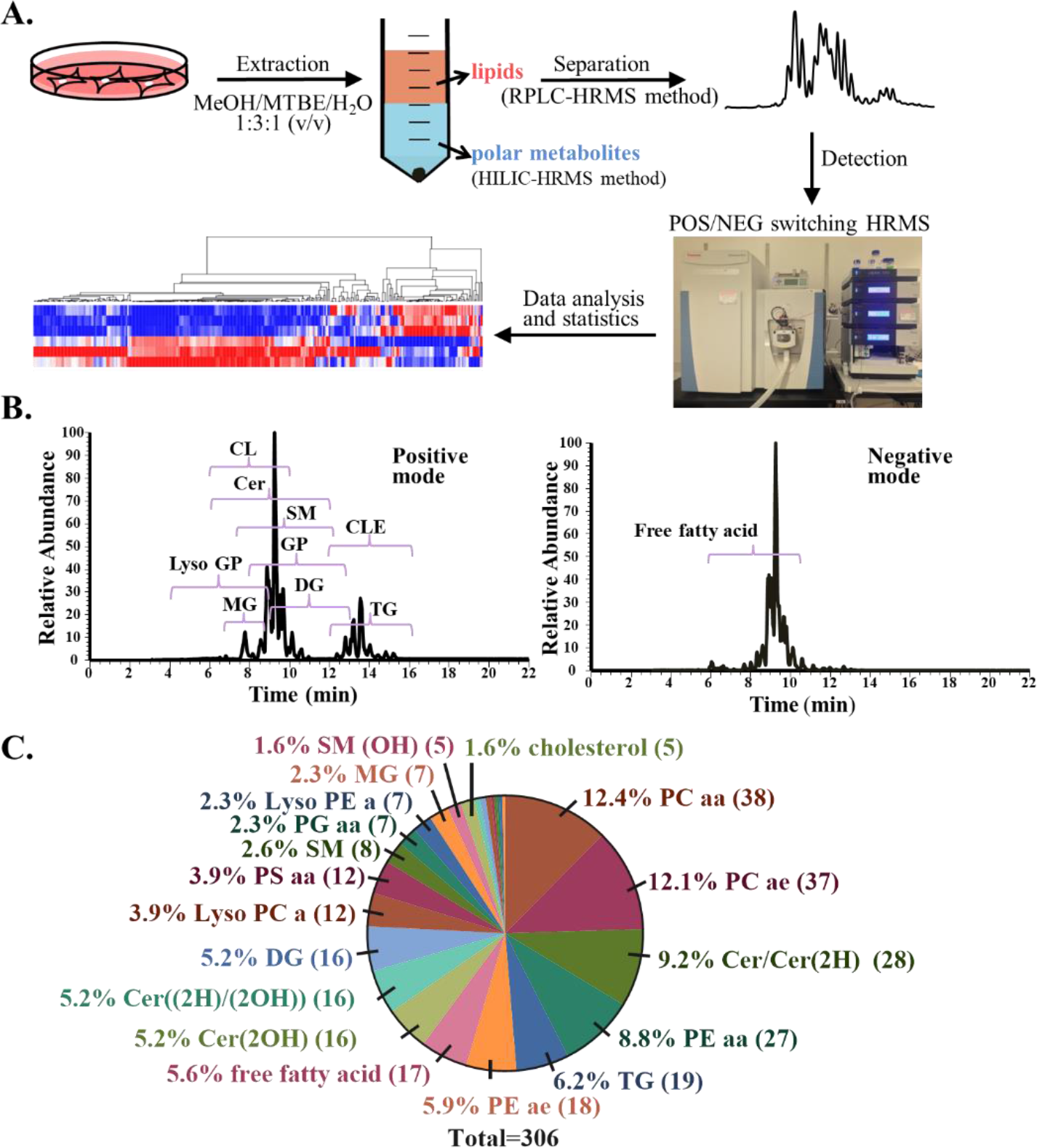
Description of lipidomics platform. (A) Workflow of lipidomics platform employing reversed phase LC coupled with high resolution MS (HRMS). (B) Total ion chromatography of lipids extracted from HCT116 using reversed phase LC coupled with HRMS operated at positive mode (left panel) and negative mode (right panel). Time denotes the retention time. (C) Lipid profile of HCT116 cells. Number in the parenthesis indicates the number of detected lipids for that lipid class. Abbreviations: CL, cholesterol; Cer, ceramides; SM, sphingomyelin; CLE, cholesterol ester; GP, glycerophospholipid; Lyso GP, lysoglycerophospholipid; MG, monoacylglyceride; DG, diacylglyceride; TG, triacylglyceride. PC aa, phosphatidylcholine (diacyl bonds). PC ae, phosphatidylcholine (acyl/ether bonds); Cer(2H), dihydroceramide; PE aa, phosphatidylethanolamine (diacyl bonds); PE ae, phosphatidylethanolamine (acyl/ethyl bonds); Cer(2OH), 2-hydroxyacyl-ceramide; Cer((2H)/(2OH)), 2-hydroxyacyl-dihyroceramide; Lyso PC a, lysophosphatidylcholine (acyl bond), PS aa, phosphatidyl serine (diacyl bonds); PG aa, phosphatidyl glycerol; Lyso PE a, lysophosphatidylethanolamine (acyl bond); SM(OH), hydroxysphingomyelin.

We next evaluated the quantitative capabilities of this platform using cultured HCT 116 cells. Lipids were first extracted from 1.12 × 10^7^ cells then followed by a serial dilution with a dilution factor of 5, resulting in 8 samples with different concentrations allowing for a span of analysis of over 4 orders of magnitude of concentration. Figure 2A presents a histogram of MS intensity values (integrated peak area within the defined retention time window for each m/z) across the targeted lipids extracted from 2.4 × 10^4^ cells. Figure 2B demonstrates a strong correlation between coefficient of variation (CV) of biological triplicates and the corresponding MS intensity (R^2^= −0.58, p = 0.04, Spearman’s correlation coefficient). Notably, smaller intensity values tend to have more interference from the background, thus resulting in a larger CV. For metabolites with MS intensities higher than 1 × 10^6^, the CV was found to be 17.9% (at 75^th^ percentile), while for MS intensities less than 1 × 10^5^, the CV varies to a larger extent (83.6% at the 75^th^ percentile). Therefore, we defined an MS intensity of 1 × 10^5^ as the noise level, and in Figure 2C, data were processed further by imputing intensities lower than 1 × 10^5^ with a value of 1 × 10^5^, as described in the methods section. The histogram of the linear regression of MS intensity is shown in Figure 2C. A linear regression analysis of concentrations (excluding the highest saturated and unsaturated 3 background concentrations) shows that ~98% of the detected metabolites have r^2^ values larger than 0.85. Thus, the integrated MS intensity can accurately represent metabolite relative levels over 3 orders of magnitude in a linear range.

**Figure 2.**
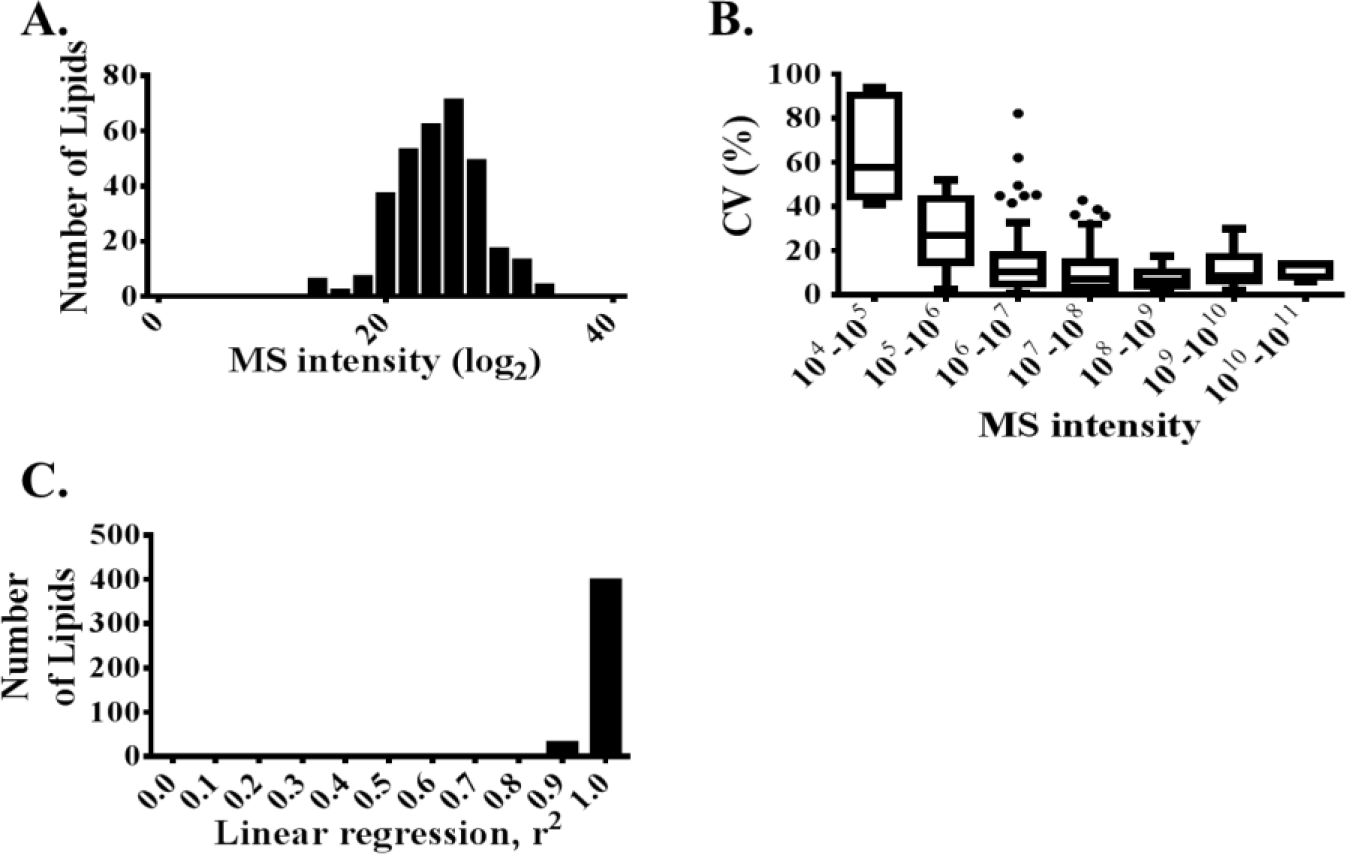
Evaluation of lipidomics platform. (A) The intensity distribution of lipids detected from 2.4 × 10^4^ of HCT116 cells. An average of n=3 biological replicates are used. (B) The relationship between coefficient of variation (CV) of triplicate samples and MS intensity. The box plot shows the 75^th^/25^th^ percentile, and the bar represents the median. (C) Linear regression analysis of the MS intensity and concentrations with over 3 orders of magnitude of each lipid detected in HCT116 cells. The number of lipids with a given r^2^ value range is shown.

A major consideration with lipidomics is the necessity to avoid contamination during sample preparation especially when organic solvents may interact with plastic tubing. Thus, we compared MeOH/MTBE/H_2_O extraction in a plastic tube, glass tube, and with 80% methanol/water extraction in glass tube as a control to evaluate MeOH/MTBE/H_2_O extraction procedure. A heatmap in Figure S1 demonstrates that MeOH/MTBE/H_2_O extraction in a plastic tube is comparable to the same extraction in a glass tube, and yields a broader coverage compared to 80% methanol/water extraction. Furthermore, our data show that interference from the background is negligible either using a plastic tube or a glass tube for MeOH/MTBE/H_2_O extraction. Figure S1 also demonstrates that the sample is relatively stable for at least 16h at 4°C after reconstitution. Together, with high resolution mass spectrometry, over 300 lipids are detected in HCT116 cells, with accurate quantitation over 3 orders of magnitude in concentration. Thus, the developed lipidomics method allows for convenient, simultaneous extraction, and detection of both lipids and polar metabolites in the same sample.

### Evaluation of the Lipidomic Response to Metformin Treatment

After developing this platform, we sought to next use it for a preliminary investigation to a question relevant to biomedical research as a case study. As discussed, the influence of metformin, a widely used diabetes drug, on lipid metabolism is not clear. Thus, the developed lipidomics method was further applied to study the response to metformin in HeyA8 OvCa cells, a model we have previously published to study the effects of metformin on polar metabolism (17). HeyA8 OvCa cells were cultured without (control group) or with (metformin group) metformin (Figure 3A). Figure 3B shows the antiproliferative effects of metformin. The generated intracellular lipid profiles from the metformin group and the control group were first analyzed by a principle component analysis (PCA), revealing metformin treatment as the major source of variation determined by its contribution to the first principal component (Figure 3C). This is further confirmed by a visual inspection of the heatmap in Figure 3D, where columns represent different lipids and rows represent biological samples. 256 lipids (isomers are not differentiated and combined together as one lipid) detected in HeyA8 OvCa cells included the following species: 35 PC aa, 29 PC ae, 26 Cer/Cer(2H), 23 PE aa, 20 TG, 18 PS aa, 16 DG, 13 PE ae, 13 free fatty acids, 10 Cer((2H)/(2OH)), 8 lyso PC a, 8 SM, 6 Cer(2OH), 5 PG aa, 5 cholesterol, 4 phosphatidylserine acyl/ether (PS ae), 4 Lyso PE a, and 4 MG (Figure 3E). Among the 256 detected lipids, 99 lipids (39%) were significantly increased (p ≤0.05, t test) and 11 lipids (4%) were significantly reduced (p ≤0.05, t test) after metformin treatment, while 146 lipids (57%) remain unchanged. The 99 significantly increased lipids (Figure 3F) include: 17 TG (Figure 3G), 16 Cer/Cer(2H), 12 PC aa, 12 PE aa, 7 free fatty acid, 7 PE ae, and 5 PC ae etc. 11 significantly decreased lipids include: 4 PC ae, 2 DG, 2 PC aa, 1 lyso PC a, 1 PE aa, 1 SM(OH). Figure 3G shows that 17 out of 20 detected TGs increased after metformin treatment. Together our data demonstrates that most cellular lipids are significantly increased upon metformin treatment. Meanwhile, the impact on different lipid classes varies from one to another, for example, even within the same lipid class PC ae, there are 5 lipids significantly increased while 4 lipids significantly decreased after metformin treatment (totally 29 detected).

**Figure 3.**
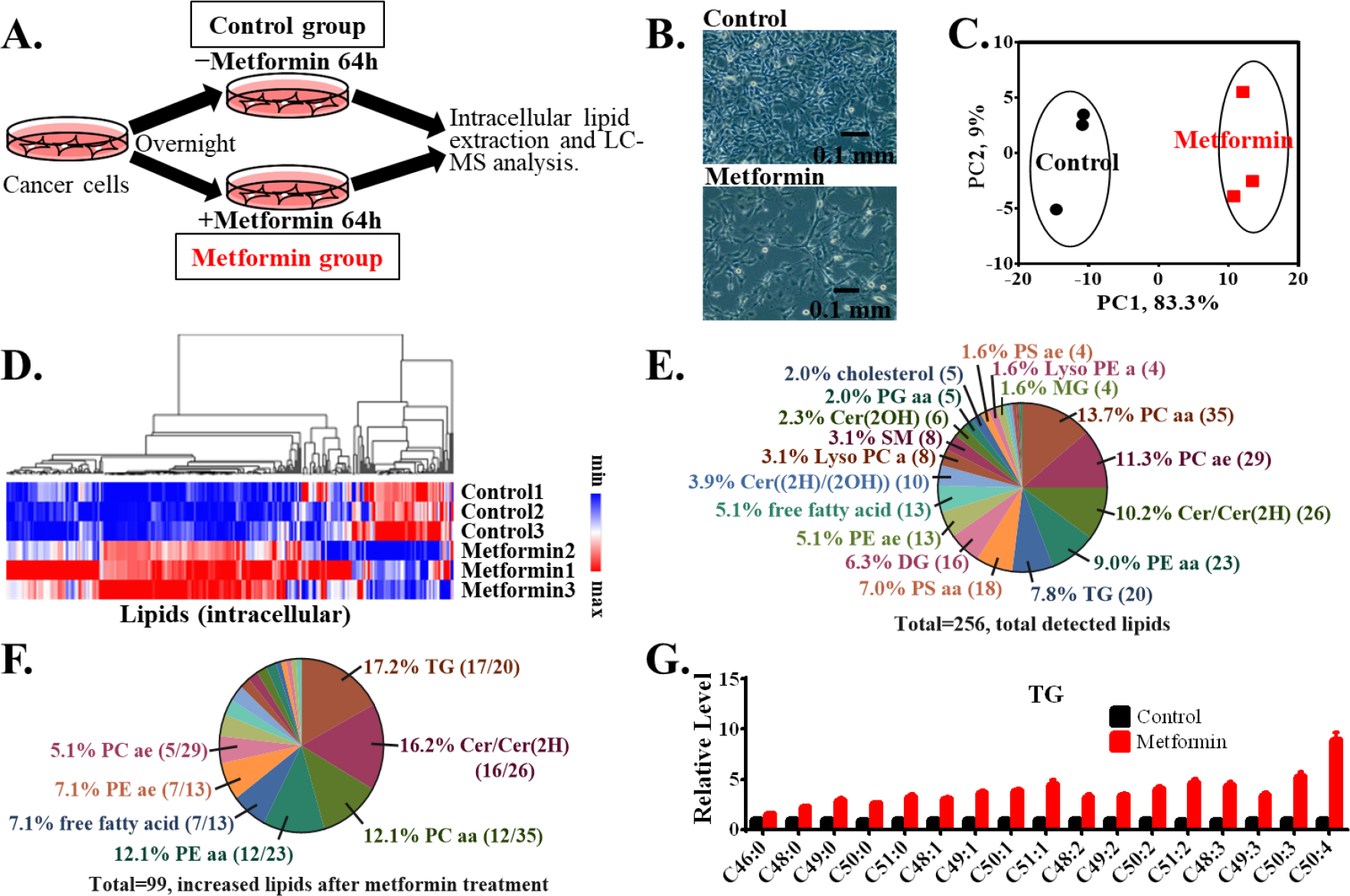
The lipid profile alterations in HeyA8 ovarian cancer cells in response to metformin treatment. (A) Schematic depicting the workflow for the lipid analysis of HeyA8 OvCa cells with or without metformin treatment. (B) Representative photos of HeyA8 OvCa cell cultured in the absence (control group) and in the presence (metformin group) of metformin for 64 hrs. Scale bar, 0.1mm. (C) Principle component analysis (PCA) of intracellular lipids in control and metformin treatment groups. (D) The heatmap of the relative level of intracellular lipids in cells with or without metformin treatment. (E) Lipid profile of HeyA8 OvCa cells. Number in the parenthesis indicates the number of detected lipids for that lipid class. (F) Summary of the composition of 99 lipids with significant increase (relative FC ≥ 0.5 & p ≤ 0.05, t test) after metformin treatment. Relative FC is defined to be the signal difference of metformin and control group divided by control group signal. The 1^st^ number in parenthesis indicates number of lipids significantly increased out of total detected lipids (the 2^nd^ number in parenthesis) for that lipid class. (G) The relative level of significantly increased 17 TGs in HeyA8 OvCa cells with or without metformin treatment.

### Alterations of Lipid Biosynthesis from Glucose in Response to Metformin Treatment

To further illustrate the utility of our platform, we studied the effects of metformin on incorporation of glucose into cellular lipids by performing ^13^C-tracing using [^13^C_6_]-glucose. In consideration of the backbone differences of these lipid classes, we report the effects of metformin treatment on glycerolipids/glycerophospholipids (glycerol backbone) and ceramides (sphingosine base backbone) separately. Alterations to glycerolipids/glycerophospholipids biosynthesis are shown in Figure 4. Figure 4A shows the workflow of the [^13^C_6_]-glucose tracing assay. Figure 4B is a scheme of incorporation of [^13^C_6_]-glucose into fatty-acyl chains (non-polar tail) and the glycerol backbone of glycerolipids/glycerophospholipids. As glycerol-3-phosphate is the precursor of the glycerol backbone and fatty-acyl chains are produced *de novo* with acetyl-CoA, we first examined their mass isotopologue patterns (Figure 4C and 4D). Due to the low signal of acetyl-CoA, acetyl-carnitine which has shown to correlate well with acetyl-coA levels (19) is used to represent acetyl-CoA. The majority of glycerol-3-phosphate and acetylcarnitine are derived from glycolysis and glycerol-3-phosphate is not affected by metformin while acetylcarnitine level decreased after metformin treatment. We then checked the incorporation of these two into lipids: TG C50:1 (Figure 4E), PE aa C38:1 (Figure 4F) and PC ae C36:5 (Figure 4G). Figures 4E-F show that d*e novo* fatty acid synthesis is inhibited while contribution to the glycerol backbone (mostly derived from glycolysis) increased after metformin treatment. Further investigation shows that relative levels of glycerol-3-phosphate mass isotopologues (Figure 4H) and total levels of glycerol-3-phosphate (Figure 4I) are not affected by metformin treatment, while the relative levels of TG C50:1 mass isotopologues increased (Figure 4J).

**Figure 4.**
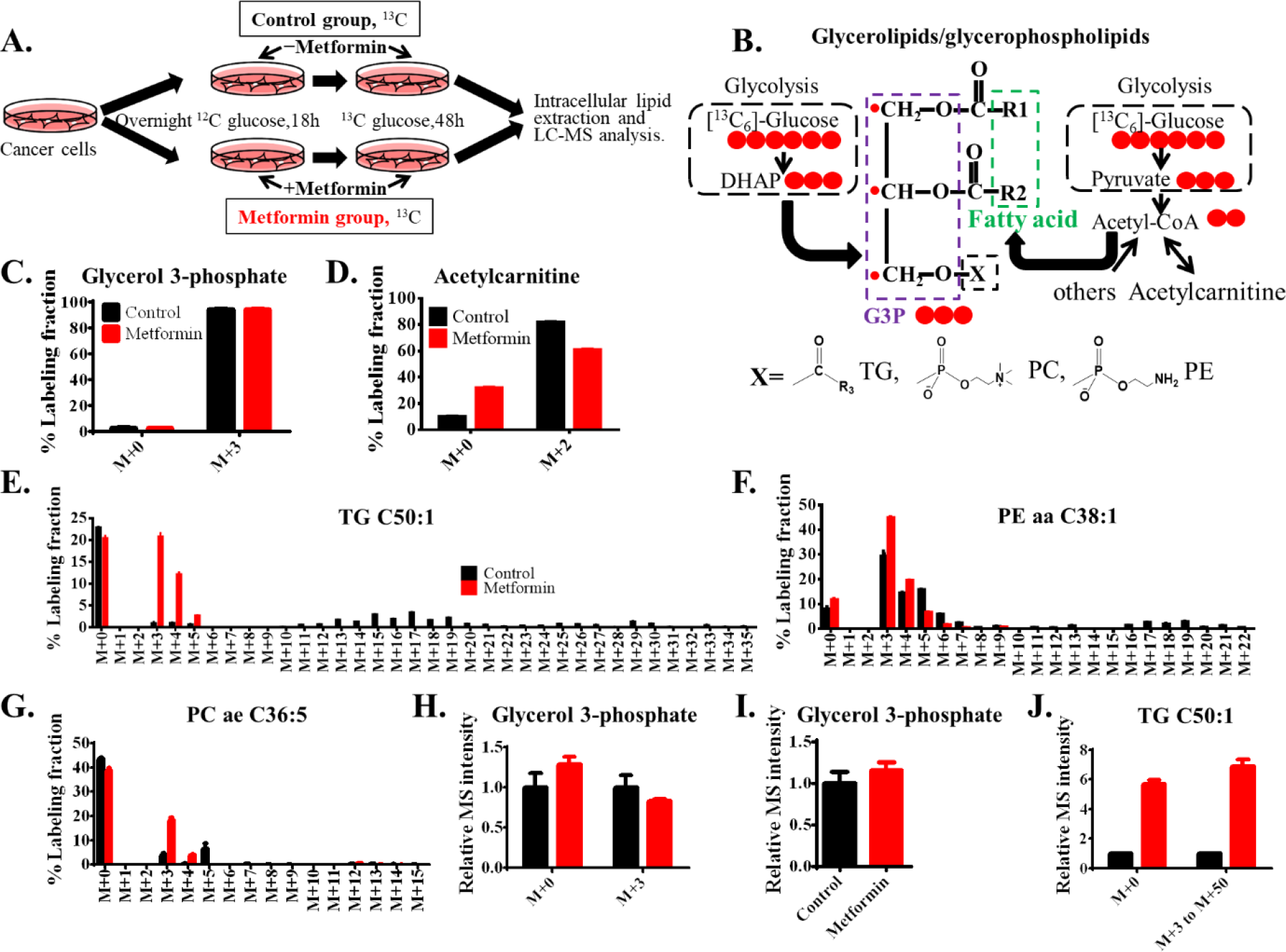
Alterations of glycerolipids and glycerophospholipids biosynthesis from glucose in cancer cells in response to metformin treatment. (A) Schematic depicting the workflow for the [^13^C_6_]-glucose tracing assay and lipid analysis in HeyA8 OvCa cells. (B) A schematic representation of [^13^C_6_]-glucose incorporation into glycerol backbones and fatty-acyl chains (non-polar tails) of glycerolipids and glycerolphospholipids. X represents the polar head and R represents non-polar tail, except that for triacylglycerides, X is a fatty-acyl chain. ^13^C isotopologue distribution of (C) Glycerol 3-phosphate, (D) Acetylcarnitine, (E) TG C50:1, (F) PE aa C38:1, (G) PC ae C36:5. The relative MS intensity of ^13^C mass isotopologues (H) and the total level (I) of glycerol 3-phosphate from cells cultured in RPMI with [^13^C_6_]-glucose. (J) The relative MS intensity of ^13^C mass isotopologues of TG C50:1 from cells cultured in RPMI with [^13^C_6_]-glucose. Natural abundance was corrected for the reported % labeling fraction.

For ceramides (Figure 5), different from glycerolipids/glycerophospholipids, serine is the precursor of the sphingosine base backbone (Figure 5A). Figure 5B shows that glucose-derived serine decreased after metformin treatment. Incorporation of serine into ceramides Cer C24:1 (Figure 5C) and Cer(2H) C24:0 (Figure 5D) increased after metformin treatment while *de novo* synthesis of fatty-acyl chain being inhibited. The relative level of serine mass isotopologues (Figure 5E) and total level of serine (Figure 5F) demonstrate that all serine species increased upon metformin treatment.

**Figure 5.**
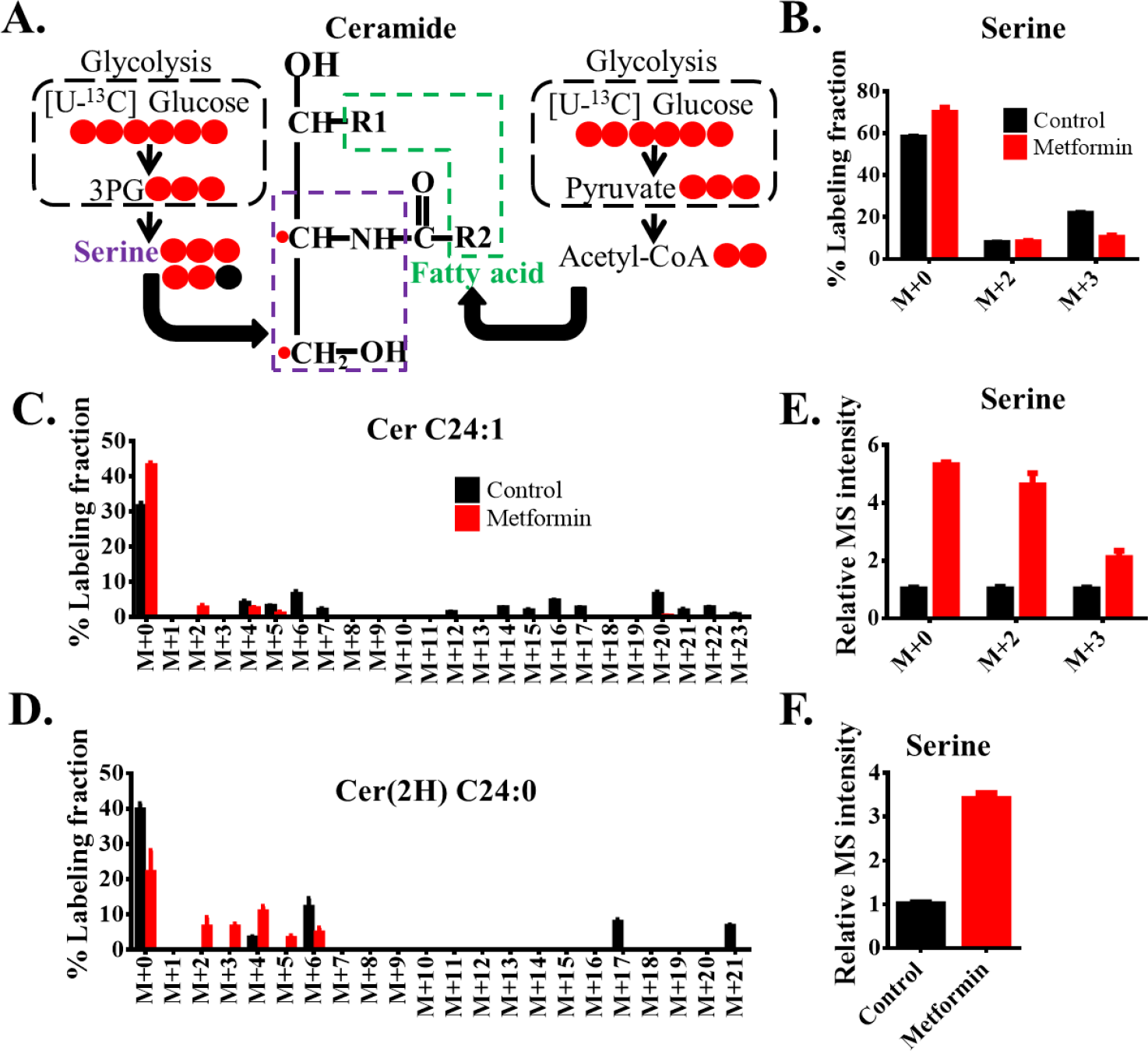
The effect of metformin on the ceramide biosynthesis from glucose in HeyA8 ovarian cancer cells. (A) A schematic representation of [^13^C_6_]-glucose incorporation into sphingosine base backbones and fatty-acyl chains of ceramides. R represents the fatty-acyl chain. ^13^C mass isotopologue distribution of serine (B), Cer C24:1 (C) and Cer(2H) C24:0 (D). The relative MS intensity of ^13^C mass isotopologues (E) and the total level (F) of serine from cells cultured in RPMI medium containing [^13^C_6_]-glucose. Natural abundance was corrected for the reported % labeling fraction.

## Discussion

In summary, we developed a metabolomics and a lipidomics platform thus that measures both lipids and polar metabolites by employing a MTBE extraction, followed by HILIC and RPLC chromatography methods coupled with HRMS. This workflow allows for measuring of both lipids and polar metabolites from a single sample in two short chromatography runs. Thus, in about one hour of experiment time, over 500 metabolites can be quantified in one sample. As a case study, we used the developed platform to study the response to metformin treatment, which preliminarily suggested metformin has differential effects on different lipid classes. In the presence of lipid-rich medium, certain lipids accumulated after metformin treatment, which is consistent with a recent study showing that the certain classes of lipids accumulated in breast cancer cells after treatment with a fatty acid synthase inhibitor (20). Further, as a proof of concept we measured with stable isotopes, the incorporation of glucose into lipids. As a limitation, these very preliminary experiments were performed in replete media containing high glucose and lipid levels and these results undoubtedly will change when these environmental variables are affected. Nevertheless, the analysis clearly demonstrates our ability to robustly profile lipidomic remodeling in the face of a cellular perturbation.

Notably, additional limitations remain despite the advances made in this study. Certain bioactive lipids such as prostaglandins are not measured using this methodology due to the relatively low abundance and low ionization efficiency of these lipids when electrospray ionization is used. In addition, the complexity of studying lipid metabolism also arises from the numerous isomers, and this current method doesn’t separate or identify many of these isomers. To address these issues, additional analysis is required. For instance, chemical derivatization, chiral chromatography, atmospheric pressure chemical ionization source and tandem mass spectrometry can each be incorporated to improve the detection limit and enhance separation (21). Separately, ion mobility mass spectrometry coupled to high resolution or tandem mass spectrometry also shows promise in separating lipid isomers (22, 23). To identify the double bond position in unsaturated lipids, ozone electrospray ionization has been employed to oxidize double bonds and introduce hydroxy groups (24), which enables the generation of characteristic product ions near the double bond to determine the double bond location. Nevertheless, our method does provide substantial information of the biological importance of different lipid classes and can be achieved using straightforward sample preparation and simple, fast chromatography.

Furthermore, the subcellular location of lipids is also important. It has been reported that the same lipid classes in different organelles play a different role in regulating cellular process (8, 25). In this study, the data represent the lipids from the whole cells, but in the future, it will be of great importance to investigate lipid remodeling in different cellular cell organelles by methods such as mass spec imaging.

## Conclusions

All together our study demonstrates that a MTBE extraction, followed by HILIC and RPLC coupled to high resolution mass spectrometry allows for a metabolomics investigation. Furthermore, this workflow can also be adapted to investigate lipid metabolism in other systems, such as cultured cells or animals subjected to treatments targeting lipid metabolism.

## Authors’ Contributions

JWL, XL, JL conceived the study, designed the experiments, and performed data analysis and interpretation. JL, XL and JWL wrote the paper. ZX helped interpreting data.

## Acknowledgment

We acknowledge support from National Institutes of Health awards R01CA193256, R00 CA16899 to JWL. We also thank members of the Locasale Lab for helpful discussions. No conflicts of interest are reported at this time.

## References

1. Yang, K., and X. Han. 2016. Lipidomics: Techniques, Applications, and Outcomes Related to Biomedical Sciences. Trends Biochem Sci 41: 954–969.

2. Brugger, B. 2014. Lipidomics: analysis of the lipid composition of cells and subcellular organelles by electrospray ionization mass spectrometry. Annu. Rev. Biochem. 83: 79–98.

3. Han, X., and R. W. Gross. 2003. Global analyses of cellular lipidomes directly from crude extracts of biological samples by ESI mass spectrometry: a bridge to lipidomics. J. Lipid Res. 44: 1071–1079.

4. Cajka, T., and O. Fiehn. 2016. Toward Merging Untargeted and Targeted Methods in Mass Spectrometry-Based Metabolomics and Lipidomics. Anal Chem 88: 524–545.

5. Liu, X., and J. W. Locasale. 2017. Metabolomics: A Primer. Trends Biochem.Sci. 42: 274–284.

6. Patti, G. J., O. Yanes, and G. Siuzdak. 2012. Metabolomics: the apogee of the omics trilogy. Nat Rev Mol Cell Bio 13: 263–269.

7. Johnson, C. H., J. Ivanisevic, and G. Siuzdak. 2016. Metabolomics: beyond biomarkers and towards mechanisms. Nat Rev Mol Cell Bio 17: 451–459.

8. Gao, X., K. Lee, M. A. Reid, S. M. Sanderson, C. P. Qiu, S. Q. Li, J. Liu, and J. W. Locasale. 2018. Serine Availability Influences Mitochondrial Dynamics and Function through Lipid Metabolism. Cell Rep 22: 3507–3520.

9. Quehenberger, O., A. M. Armando, and E. A. Dennis. 2011. High sensitivity quantitative lipidomics analysis of fatty acids in biological samples by gas chromatography-mass spectrometry. Biochim. Biophys. Acta 1811: 648–656.

10. Dennis, E. A., R. A. Deems, R. Harkewicz, O. Quehenberger, H. A. Brown, S. B. Milne, D. S. Myers, C. K. Glass, G. Hardiman, D. Reichart, A. H. Merrill, M. C. Sullards, E. Wang, R. C. Murphy, C. R. H. Raetz, T. A. Garrett, Z. Q. Guan, A. C. Ryan, D. W. Russell, J. G. McDonald, B. M. Thompson, W. A. Shaw, M. Sud, Y. H. Zhao, S. Gupta, M. R. Maurya, E. Fahy, and S. Subramaniam. 2010. A Mouse Macrophage Lipidome. J Biol Chem 285: 39976–39985.

11. Fauland, A., H. Kofeler, M. Trotzmuller, A. Knopf, J. Hartler, A. Eberl, C. Chitraju, E. Lankmayr, and F. Spener. 2011. A comprehensive method for lipid profiling by liquid chromatography-ion cyclotron resonance mass spectrometry. J Lipid Res 52: 2314–2322.

12. Yore, M. M., I. Syed, P. M. Moraes-Vieira, T. Zhang, M. A. Herman, E. A. Homan, R. T. Patel, J. Lee, S. Chen, O. D. Peroni, A. S. Dhaneshwar, A. Hammarstedt, U. Smith, T. E. McGraw, A. Saghatelian, and B. B. Kahn. 2014. Discovery of a class of endogenous mammalian lipids with anti-diabetic and anti-inflammatory effects. Cell 159: 318–332.

13. Matyash, V., G. Liebisch, T. V. Kurzchalia, A. Shevchenko, and D. Schwudke. 2008. Lipid extraction by methyl-tert-butyl ether for high-throughput lipidomics. J Lipid Res 49: 1137–1146.

14. t’kindt, R., E. D. Telenga, L. Jorge, A. J. M. Van Oosterhout, P. Sandra, N. H. T. Ten Hacken, and K. Sandra. 2015. Profiling over 1500 Lipids in Induced Lung Sputum and the Implications in Studying Lung Diseases. Anal Chem 87: 4957–4964.

15. Breitkopf, S. B., S. J. H. Ricoult, M. Yuan, Y. Xu, D. A. Peake, B. D. Manning, and J. M. Asara. 2017. A relative quantitative positive/negative ion switching method for untargeted lipidomics via high resolution LC-MS/MS from any biological source. Metabolomics 13: 21.

16. Liu, X., Z. Ser, and J. W. Locasale. 2014. Development and Quantitative Evaluation of a High-Resolution Metabolomics Technology. Anal. Chem. 86: 2175–2184.

17. Liu, X., I. L. Romero, L. M. Litchfield, E. Lengyel, and J. W. Locasale. 2016. Metformin Targets Central Carbon Metabolism and Reveals Mitochondrial Requirements in Human Cancers. Cell Metab. 24: 728–739.

18. Wheaton, W. W., S. E. Weinberg, R. B. Hamanaka, S. Soberanes, L. B. Sullivan, E. Anso, A. Glasauer, E. Dufour, G. M. Mutlu, G. R S. Budigner, and N. S. Chandel. 2014. Metformin inhibits mitochondrial complex I of cancer cells to reduce tumorigenesis. eLife 3: 18.

19. Liu, X., S. Sadhukhan, S. Y. Sun, G. R. Wagner, M. D. Hirschey, L. Qi, H. N. Lin, and J. W. Locasale. 2015. High-Resolution Metabolomics with Acyl-CoA Profiling Reveals Widespread Remodeling in Response to Diet. Mol. Cell. Proteomics 14: 1489–1500.

20. Alwarawrah, Y., P. Hughes, D. Loiselle, D. A. Carlson, D. B. Darr, J. L. Jordan, J. Xiong, L. M. Hunter, L. G. Dubois, J. W. Thompson, M. M. Kulkarni, A. N. Ratcliff, J. J. Kwiek, and T. A. J. Haystead. 2016. Fasnall, a Selective FASN Inhibitor, Shows Potent Anti-tumor Activity in the MMTV-Neu Model of HER2(+) Breast Cancer. Cell Chem Biol 23: 678–688.

21. Lee, S. H., M. V. Williams, R. N. DuBois, and I. A. Blair. 2003. Targeted lipidomics using electron capture atmospheric pressure chemical ionization mass spectrometry. Rapid Commun Mass Sp 17: 2168–2176.

22. Kyle, J. E., X. Zhang, K. K. Weitz, M. E. Monroe, Y. M. Ibrahim, R. J. Moore, J. Cha, X. Sun, E. S. Lovelace, J. Wagoner, S. J. Polyak, T. O. Metz, S. K. Dey, R. D. Smith, K. E. Burnum-Johnson, and E. S. Baker. 2016. Uncovering biologically significant lipid isomers with liquid chromatography, ion mobility spectrometry and mass spectrometry. Analyst 141: 1649–1659.

23. Groessl, M., S. Graf, and R. Knochenmuss. 2015. High resolution ion mobility-mass spectrometry for separation and identification of isomeric lipids. Analyst 140: 6904–6911.

24. Thomas, M. C., T. W. Mitchell, and S. J. Blanksby. 2006. Ozonolysis of phospholipid double bonds during electrospray ionization: A new tool for structure determination. J Am Chem Soc 128: 58–59.

25. Yang, Y., M. Lee, and G. D. Fairn. 2018. Phospholipid subcellular localization and dynamics. J. Biol. Chem. 293: 6230–6240.

