## Supplemental Figure 1, Supplemental Table 1-2 for "Quantitative evaluation of a high resolution lipidomics platform"

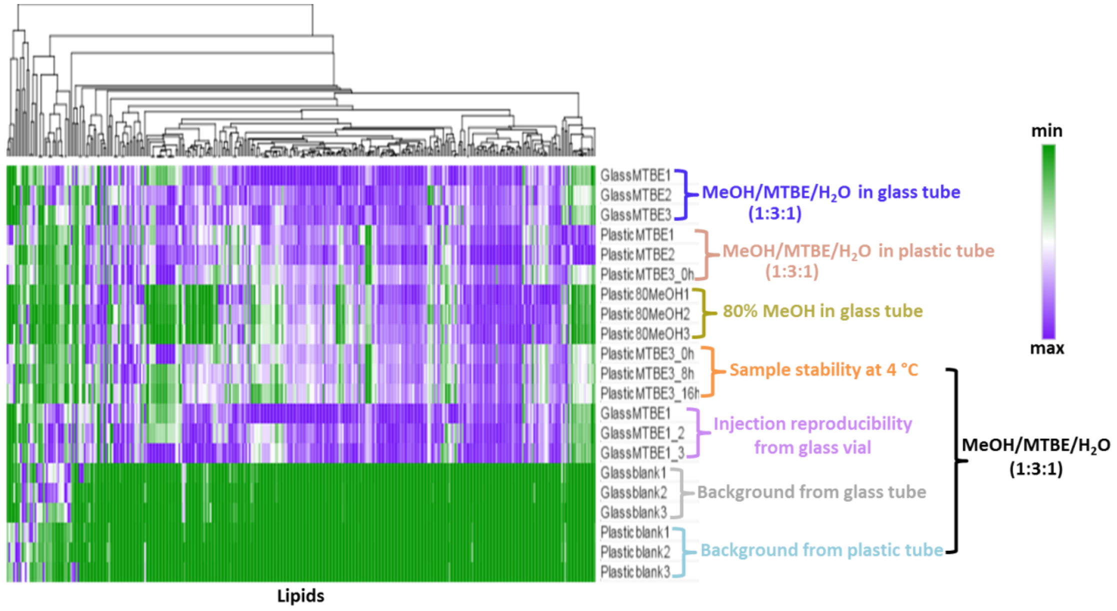


**Supplementary Figure S1.** Evaluation of MeOH/MTBE/$H_2O$ (1:3:1) extraction with plastic tubing. Comparison of MeOH/MTBE/$H_2O$ (1:3:1) extraction in a glass tube, in a plastic tube and 80% MeOH in a glass tube. Sample stability at 4$^o$C over 16h and injection reproducibility from a glass vial using MeOH/MTBE/$H_2O$ (1:3:1) extraction. Comparison of background from MeOH/MTBE/$H_2O$ (1:3:1) extraction in a glass and a plastic tube.

**Supplementary Table S1 Adduct and detection ion mode for lipids**

| **Lipid Class** | **Adduct** | **Ionization mode** |
|---|---|---|
| Free fatty acid | $H^+$ | Negative |
| Cholesterol | $H^+$ (waterloss) | Positive |
| Ceramides | $H^+$ | Positive |
| Sphingomyelin | $H^+$ | Positive |
| Cholesterol ester | $H^+$ | Positive |
| Glycerophospholipid | $H^+$ | Positive |
| Lyso glycerophospholipid | $H^+$ | Positive |
| Monoacylglyceride | $H^+$ | Positive |
| Diacylglyceride | $NH^{4+}$ | Positive |
| Triacylglyceride | $NH^{4+}$ | Positive |

**Supplementary Table S2 Abbreviations of lipids**

| **Abbreviation** | **Full biochemical name** |
| --- | --- |
| CL | Cholesterol |
| CLE | Cholesterol ester |
| Cer | Ceramide |
| Cer(2H) | dihydroceramide |
| Cer(2OH) | 2-hydroxyacyl-ceramide |
| Cer((2H)/(2OH)) | 2-hydroxyacyl-dihyroceramide |
| GP | Glycerophospholipid |
| PC aa | Phosphatidylcholine diacyl |
| PC ae | Phosphatidylcholine acyl-ether |
| PE aa | Phosphatidylethanolamine diacyl |
| PE ae | Phosphatidylethanolamine acyl-ethyl |
| PS aa | Phosphatidylserine diacyl |
| PS ae | phosphatidylserine acyl-ether |
| PG aa | Phosphatidyl glycerol diacyl |
| Lyso GP | Lyso glycerophospholipid |
| Lyso PC a | Lysophosphatidylcholine acyl |
| Lyso PE a | Lysophosphatidylethanolamine acyl |
| MG | Monoacylglyceride |
| DG | Diacylglyceride |
| TG | Triacylglyceride |
| SM | Sphingomyelin |
| SM(OH) | Hydroxysphingomyelin |
